## Supplementary material for "The Balance between B55α and Greatwall expression levels predicts sensitivity to Greatwall inhibition in cancer cells": A detailed methodology for mass spectrometry can be found 882 in the Supplementary material.

**Sample preparation for phosphoproteomics analysis.**

Sample preparation for mass spectrometry analysis was performed as described previously [1, 2]. Cell pellets were lysed with a Urea buffer (8M urea in 20mM in HEPES, pH 8.0 supplemented with 1mM $Na_3VO_4$, 1mM NaF, 1mM $Na_2H_2P_2O_7$ and 1mM sodium β-glycerophosphate), sonicated for 30 cycles (30 s on 30 s off) in a Diagenode Bioruptor® Plus and insoluble material was removed by centrifugation (13,000 rpm for 10 min at 4°C). Protein concentration was determined using BCA Protein Assay Kit. 110 µg of extracted proteins in a final volume of 200 uL were reduced with dithiothreitol (DTT, 10 mM) for 1 h at 25 °C, and alkylated with Iodoacetamide (IAM, 16.6 mM) for 30 min at 25 °C. Then, samples were diluted with 20 mM HEPES (pH 8.0) to a final concentration of 2 M urea and digested with equilibrated trypsin beads (50% slurry of TLCK-trypsin) overnight at 37 °C. The equilibration of the beads was performed by washes with 20 mM HEPES; pH 8.0. After digestion and trypsin bead removal by centrifugation (2000×g for 5 min at 5 °C), peptide solutions were transferred into 96 well plates and acidified by adding TFA to a final concentration of 0.1%. Then, samples were desalted and subjected to phosphoenrichment using the AssayMAP Bravo (Agilent Technologies) platform. For desalting, protocol peptide clean-up v3.0 was used. Reverse phase S cartridges (Agilent, 5 µL bed volume) were primed with 250 µL 99.9% acetonitrile (ACN) with 0.1%TFA and equilibrated with 250 µL of 0.1% TFA at a flow rate of 10 µL/min. The samples were loaded (770 µL) at 20 µL/min, followed by an internal cartridge wash with 250 µL of 0.1% TFA at a flow rate of 10 µL/min. Peptides were then eluted with 55 µL of 70% ACN, 0.1% TFA into 96 protein LoBind plates containing 50 uL of 1M Glycolic Acid with 50% ACN, 5%TFA. Following the Phospho Enrichment v 2.1 protocol, phosphopeptides were enriched using 5µl Assay MAP $TiO_2$ cartridges on the Assay MAP Bravo platform. The cartridges were primed with 100µl of 5% ammonia solution with 15% ACN at a flow rate of 300 µL/min and equilibrated with 50 µL loading buffer (1M glycolic acid with 80% ACN, 5% TFA) at 10 µL/min. Samples eluted from the desalting were loaded onto the cartridge at 3 µL/min. The cartridges were washed with 50 µL of loading buffer and phosphopeptides were eluted with 25 µL 5% ammonia solution with 15% ACN directly into 25 µL 10% formic acid. Phosphopeptides were lyophilized in a vacuum concentrator and stored at -80˚C.

**LC-MS/MS Analysis**

Phosphopeptides were re-suspended in 8 µL of reconstitution buffer and sonicated for 2 minutes at RT. After centrifugation (13000 rpm, 5 min, 4 °C), 5µl was loaded onto a LC-MS/MS system. This consisted of a nano flow ultra-high pressure liquid chromatography system UltiMate 3000 RSLC nano (Dionex) coupled to a Q Exactive Plus using an EASY-Spray system. The LC system used mobile phases A (3% ACN: 0.1% FA) and B (100% ACN; 0.1% FA). Peptides were loaded onto a µ-pre-column and separated in an analytical column. The gradient: 1% B for 5 min, 1% B to 35% B over 90min, following elution the column was washed with 85% B for 7 min, and equilibrated with 3% B for 7min, flow rate of 0.25 µL/min. Peptides were nebulized into the online connected Q-Exactive Plus system operating with a 2.1s duty cycle. Acquisition of full scan survey spectra (m/z 375-1,500) with a 70,000 FWHM resolution was followed by data-dependent acquisition in which the 15 most intense ions were selected for HCD (higher energy collisional dissociation) and MS/MS scanning (200-2,000 m/z) with a resolution of 17,500 FWHM. A 30s dynamic exclusion period was enabled with an exclusion list with 10ppm mass window. Overall duty cycle generated chromatographic peaks of approximately 30s at the base, which allowed the construction of extracted ion chromatograms (XICs) with at least ten data points.

**Phosphopeptides identification and quantification**

Peptide identification from MS data was automated using a Mascot Daemon (v2.8.0.1) workflow in which Mascot Distiller generated peak list files (MGF) from RAW data, and the Mascot search engine matched the MS/MS data stored in the MGF files to peptides using the SwissProt Database restricted to *Homo sapiens* (SwissProt_2021_02.fasta). Searches had an FDR of ~1% and allowed 2 trypsin missed cleavages, mass tolerance of ±10 ppm for the MS scans and ±25 mmu for the MS/MS scans, carbamidomethyl Cys as a fixed modification and oxidation of Met, PyroGlu on N-terminal Gln and phosphorylation on Ser, Thr, and Tyr as variable modifications. Pescal was used for label free quantification of the identified peptides as described before [1, 3]. The software constructed XICs for all the peptides identified in at least one of the LC-MS/MS runs across all samples. XIC mass and retention time windows were ±7 ppm and ±2 min, respectively. Quantification of peptides was achieved by measuring the area under the peak of the XICs. Individual peptide intensity values in each sample were normalized to the sum of the intensity values of all the peptides quantified in that sample. Finally, the phosphoproteomics data was processed and analysed using a public bioinformatic pipeline developed in a R environment (https://github.com/CutillasLab/protools2/). The normalised data was centered, log2 scaled and 0 values were inputted using the minimum feature value in the sample minus one. Then, statistical differences (p-values) were calculated using LIMMA [4], and then adjusted for FDR using the Benjamini-Hochberg procedure. Differences were considered statistically significant when p-values <0.05 and FDR<0.1.
